# CavitySpace: A database of potential ligand binding sites in the human proteome

**DOI:** 10.1101/2022.01.25.477691

**Authors:** Shiwei Wang, Haoyu Lin, Zhixian Huang, Yufeng He, Xiaobing Deng, Youjun Xu, Jianfeng Pei, Luhua Lai

## Abstract

The ligand binding sites of a protein provide useful information to uncover its functions and to direct the structure-based drug design. However, as binding site detection relies on the three-dimensional (3D) structural data of proteins, functional analysis based on protein ligand binding sites is formidable for proteins without structural information. Recent developments in protein structure prediction and the 3D structures built by AlphaFold provide an unprecedented opportunity for analyzing ligand binding sites in human proteins. We have used the reliable ligand binding site detection program CAVITY to analyse all the proteins in the human proteome and constructed the CavitySpace database, which is the first pocket library for predicted protein structures. CavitySpace can be used to predict protein function based on pocket information, to identify new druggable protein targets for drug design, and to search for new binding sites for known drugs for drug repurposing. CavitySpace is freely available at http://www.pkumdl.cn:8000/cavityspace/.

## Introduction

Protein-ligand interactions govern many biological processes. The specific ligand binding site (LBS) in a protein is essential for understanding its biological function and for structure-based drug design [1]. LBSs can be directly obtained from known protein-ligand complex structures. The Protein Data Bank (PDB) [2] provides the primary source of protein three-dimensional (3D) structures experimentally resolved. However, among the more than 190,000 experimentally-determined structures by 2021, only a small part were solved with bound ligand. In order to fill the gap between structures and binding sites, computational methods predicting LBSs from protein 3D structures have been developed [3]. Several pocket databases have been constructed (Table S1 gives a list of known pocket databases).

However, currently available pocket databases are limited to known protein structures. Only about 37% of human proteins have the corresponding PDB entries [4]. The protein structure prediction approaches have made great progress in the past several decades [5–7]. In 2021, AlphaFold, a deep neural network-based method developed by DeepMind has made a major breakthrough and produced protein structures with atomic accuracy even where no similar structure is known [8]. AlphaFold was then applied to build protein structure models for human proteome [9, 10], which dramatically expanded the structural coverage of human proteins.

In this work, we analysed potential ligand binding sites in the human protein structures predicted by AlphaFold and constructed a comprehensive ligand binding site database, CavitySpace. CavitySpace expands the ligand binding site space from known protein structures to predicted structures and provides a resource for protein function study and drug design.

## Material and Methods

We applied our CAVITY tool [11] to detect potential ligand binding sites from AlphaFold predicted protein structures. We also constructed a hrefPDB dataset by screening all the representative human protein structures from PDB and detected cavities for these structures. We have demonstrated that out CAVITY tool can correctly identify known binding sites from experimental or predicted protein structures. Please see the Supplementary Data for details.

### Database Introduction

#### Cavity library for AlphaFold structures

The cavity detection procedure found 237,872 cavities for the 18,672 AlphaFold structures. The druggability of each cavity was labeled as strong, medium or weak by CAVITY. Among the AlphaFold cavities, 16.3% were predicted as strong druggable cavities (Figure S1A). We further analysed the structure reliability of the residues in AlphaFold cavities. In AlphaFold structures, the predicted Local Distance Difference Test (pLDDT) was given for every residue to measure the local accuracy [8]. Structure regions with pLDDT > 90 are considered as highly reliable. Based on the pLDDT scores, we defined the ratio of the number of high confident residues (pLDDT > 90) to the total number of residues in the cavity as an Index to evaluate the reliability of the cavity structure. Among the strong AlphaFold cavities, 25.9% (Index > 0.6) contain residues with reliable structures (Figure S2).

#### Applications of the cavity library

For the 63.6% of the AlphaFold predicted human protein structures with no experimental structure information, CAVITY detected 145,444 cavities and 17.4% of them are strong druggable cavities.

As similar binding sites may bind the same or similar ligands and have similar functions, we used PocketMatch [12] to compare binding sites and the PMSmax score to evaluate the overall pocket similarity (see the Supplementary Data for details). To get meaningful results, we only analyzed the 60,913 high-quality cavities, each of which contains at least 80% residues with pLDDT > 90. These high-quality cavities, together with 50,514 hrefPDB cavities, were used to perform an all-to-all pocket comparison. We then clustered the 111,427 cavities with the Butina algorithm [13] (see Supplementary Data). With the threshold of PMSmax 0.6, 11,221 cavities did not have any similar cavities, which may be novel ligand binding sites. The other 100,206 cavities were grouped into 8,016 clusters and 538 of them contain more than 10 members. The clusters that contain known ligand binding sites can be used to study the function of proteins that contain similar cavities or to find new targets for a known ligand. For example, the crystal structure of human cysteinyl leukotriene receptor 1 binding with its antagonist zafirlukast, an FDA approved drug for asthma treatment, has been solved [14]. We selected seven top-scoring AlphaFold cavities of proteins without known PDB structures from the corresponding cavity cluster that the zafirlukast binding site belongs to (see Supplementary Data for details). Docking study showed that zafirlukast can potentially bind to these cavities with high binding affinity (Table S2).

The CavitySpace database can be used for various purposes, including identifying new druggable protein targets for drug design, predicting protein function based on pocket comparison, searching for new binding sites for known drugs for drug repurpose study, etc. It should be noted that the AlphaFold structures are currently single-chain structures, while many proteins form oligomers to be functional. We recommend that based on the CavitySpace results, users carry out further analysis of the potential binding sites with more accurate structures after carefully considering inter-domain orientations and oligomeric states using our CavityPlus webserver [15] or other cavity analysis tools.

#### The webserver

We developed the CavitySpace webserver for public usage. Users can conveniently query the database with protein name, UniProt ID or gene name and obtain the cavity details for each structure visually. All data in the cavity library can be downloaded from the CavitySpace webserver, including the strong druggable cavities, the cavity clustering results and so on. It is freely available at http://www.pkumdl.cn:8000/cavityspace/.

## Acknowledgements

We thank all the members in the Lai group for their helpful discussions and testing of the CavitySpace database. We thank the high-performance computing platform of the Peking-Tsinghua Center for Life Sciences, Peking University for providing the computational resources.

## Funding

This work has been supported in part by the Ministry of Science and Technology of China (2016YFA0502303) and the National Natural Science Foundation of China (22033001 and 21673010).

## Supplementary Data

### Data collection

All data used to construct CavitySpace were obtained from public databases. The human protein structures predicted by AlphaFold were downloaded from the AlphaFold Protein Structure Database (https://alphafold.ebi.ac.uk/). Only structures for Homo sapiens were downloaded, which contain 23391 predicted structures of 20504 sequence entries.

About 37% of human proteins can be mapped to PDB entries. Detecting LBSs from the known structures is obviously a better choice. We queried the UniProtKB database (https://www.uniprot.org/) with the UniProt ID of each sequence to retrieve UniProt entries with known PDB structures and obtained a total of 7245 UniProt records. For each UniProt entry, the structure with the best resolution was selected as a representative structure. Sometimes several PDB structures for one protein cover different domains of the same sequence. In these cases, we selected representative structures for each domain of the sequence to cover the whole protein sequence as long as possible. In addition, all PDB structures not resolved by X-ray crystallography or with resolution larger than 3.5 Å were excluded. Finally, we obtained 6967 PDB structures of 5731 UniProt entries, forming the known human PDB structure dataset (hrefPDB). All the structure files were downloaded from RCSB PDB (https://www.rcsb.org/). Because each AlphaFold structure has only one single chain, we extracted one chain from each of the known structures to keep the consistency.

### Cavity detection

We applied the CAVITY tool developed by our lab to detect all the potential cavities on protein surfaces [11]. For all the 23391 AlphaFold structures, CAVITY successfully processed 18820 (80.5%) structures. The remaining 4571 structures that CAVITY could not finish the job within a reasonable time were mainly complicated structures with relatively long protein sequences and many irregular loops. For the hrefPDB dataset containing known PDB structures, all the 6967 structures can be processed by CAVITY and 86.9% (6051) of them have at least one cavity.

### The quality of AlphaFold cavities

One important question is how different is the hrefPDB structures from the AlphaFold predicted structures for cavity detection. Thus, we performed cavity detection process for all the hrefPDB structures, producing 50,514 PDB cavities. To make a fair comparison, we extracted the subset of AlphaFold structures sharing the same UniProt IDs to the hrefPDB structures and then collected their cavities, obtaining 65,580 AlphaFold cavities. The number of cavities from the hrefPDB is smaller because part of the PDB structures is not full-sequence structure. In addition, some AlphaFold cavities locate on low confident protein regions. One of our primary concerns is finding potential bindings sites from the cavity library, so we further checked if the true ligand sites are correctly identified by CAVITY from AlphaFold structures and hrefPDB structures. The true ligand sites were defined as residues within 4 Å around bound ligands. From the hrefPDB dataset, we selected 2,439 true ligand binding sites as a test set. We found that 81% of true binding sites can be recovered from hrefPDB structures when a cavity with at least 50% residues of the true binding sites was considered as the same binding site. The number is 80% for AlphaFold structures. Such results demonstrated that our CAVITY program can successfully discover most of the true binding sites from protein structures and the AlphaFold structures are as reliable as experimentally resolved structures to be used to find potential ligand.

### Pocket comparison

we used PocketMatch to compare binding sites for function analysis [12]. PocketMatch represents each binding site as 90 lists of sorted distances capturing the shape and chemical nature of the site and then aligns them incrementally to obtain a similarity score called PMScore, which is scaled between 0 and 1, where 1 indicates identity. PocketMatch provides two type scores, one score called PMSmax implying significant similarity in the whole site and the other score called PMSmin reflecting a local sub-structural match. We select the PMSmax to evaluate the pocket similarity because it is believed to indicate biologically meaningful similarities.

### Clustering

we clustered the total 111427 cavities with the Butina algorithm [13]. We have tried different thresholds of PMSmax. With the threshold of PMSmax ≥ 0.8, 89564 cavities have no similar cavity, reminding that the threshold is too strict. With the threshold of PMSmax ≥ 0.7, 50513 cavities still have no similar cavity. The remaining cavities were grouped into 12943 clusters and 589 of them contain more than 10 cavities. When the threshold of PMSmax ≥ 0.6 was used, 11221 cavities have no similar cavity. The remaining cavities were grouped into 8016 clusters and 538 of them contain more than 10 cavities. When the threshold of PMSmax ≥ 0.5 was used, 980 cavities have no similar cavity and the remaining cavities were grouped into 230 clusters. However, the first cluster contains 31.6% of the cavities. It is obvious that the cavities cannot be classified well. Finally, we select the threshold of PMSmax ≥ 0.6 to make a clustering analysis.

### Pocket analysis

Cysteinyl leukotriene receptor 1 (CysLT1R) is a G protein-coupled receptor as well as a key player in allergic and inflammatory disorders and zafirlukast is a selective antagonist of CysLT1R [14]. In order to find potential new binding sites for zafirlukast, we investigated the cavity cluster that the zafirlukast binding site belongs to and screened all cavities with PMSmax > 0.8 that have strong druggability and do not have known PDB structures. Among 16 compliant AlphaFold cavities, we chose only one representative cavity for those cavities that were in the same domain or motif, such as the seven-transmembrane domain of GPCR and kelch motif. In addition, we abandoned cavities from Cytochrome P450. At last, we obtained 7 representative AlphaFold cavities. We performed molecular docking between target proteins and zafirlukast using AutoDock Vina 1.2 [16] (Table S2). Docking study showed that zafirlukast can bind to these cavities with high affinity, which can be experimentally tested in the future.

---

**Figure S1.**
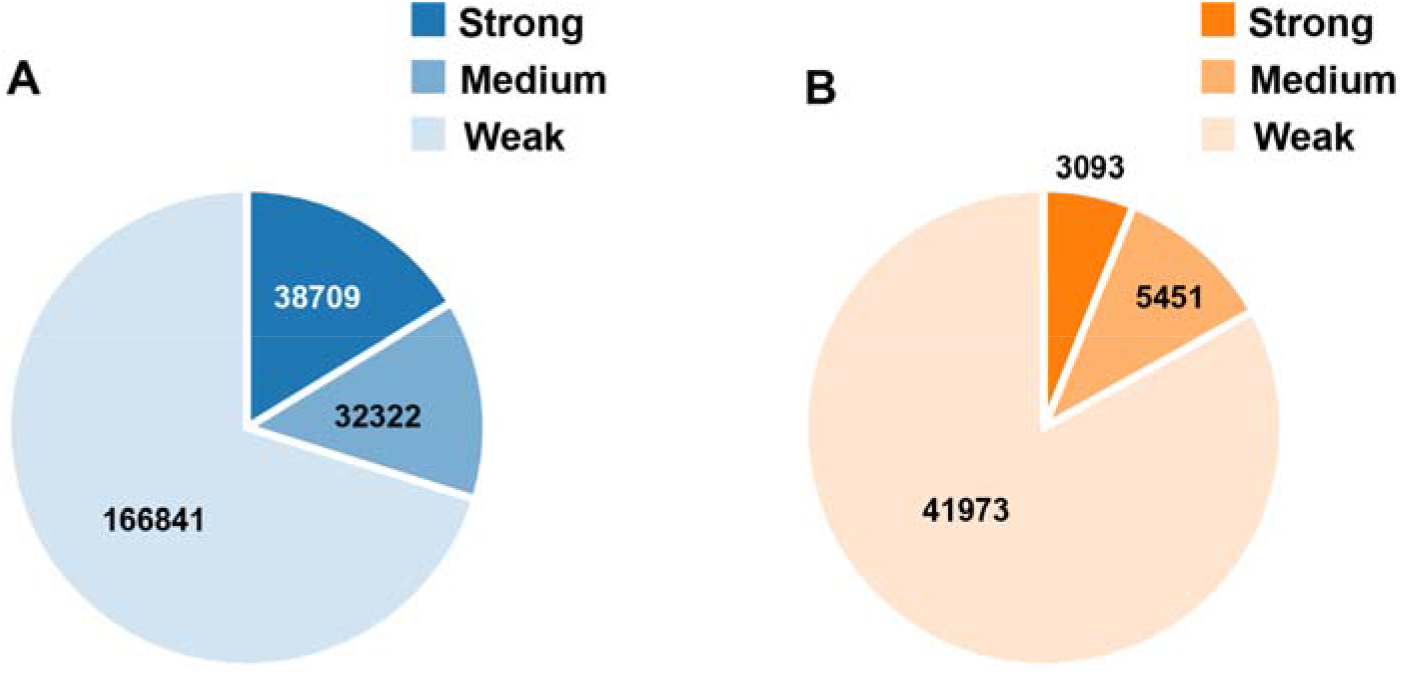
The druggability distribution of AlphaFold cavities (A) and hrefPDB cavities (B).

**Figure S2.**
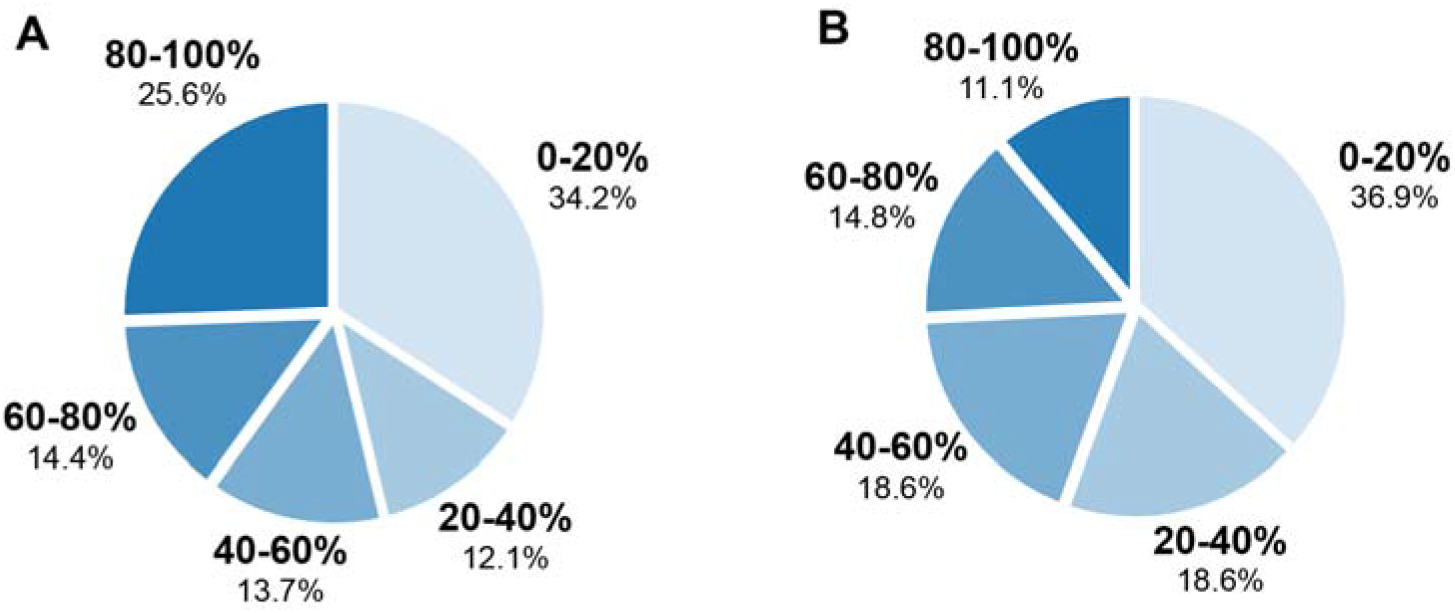
The distributions of the percentage of cavity residues with high confidence (pLDDT > 90). (A) for all cavities from AlphaFold structures and (B) for only strong druggable cavities from AlphaFold structures.

**Table S1.** Databases of protein pockets since 2004.

| Database | Publication Year | Website | Reference |
| --- | --- | --- | --- |
| ProBiS-Dock Database | 2021 | predicted small molecule and cofactor binding sites | [17] |
| HKPocket | 2019 | predicted human kinase pocket | [18] |
| PocketDB | 2018 | predicted small-molecule binding pockets | [19] |
| TuberQ | 2014 | *Mycobacterium tuberculosis* protein druggability | [20] |
| sc-PDB | 2014 | predicted ligandable binding sites | [21] |
| KLIFS | 2014 | kinase–ligand interaction fingerprints and structure | [22] |
| Bival-bind | 2014 | protein complexes with multivalent binding ability | [23] |
| FireDB | 2013 | catalytic and biologically relevant small ligand-binding residues from PDB | [24] |
| Pocketome | 2011 | experimentally solved conformational ensembles of druggable binding sites in proteins | [25] |
| PoSSuM | 2011 | similar protein–ligand binding and putative pockets | [26] |
| fPOP | 2009 | protein functional surfaces identified by analyzing the shapes of binding sites in both holo and apo forms | [27] |
| CREDO | 2009 | protein–ligand interactions with structural interaction fingerprints and novel features | [28] |
| SuperSite | 2008 | metabolite and drug binding sites in proteins | [29] |
| LigASite | 2007 | biologically relevant binding sites in proteins with known apo-structures | [30] |
| SitesBase | 2006 | structure-based protein–ligand binding site comparisons | [31] |
| PDBSite | 2005 | protein active sites and their spatial environment | [32] |
| Het-PDB Navi2 | 2004 | protein–small molecule interactions | [33] |

**Table S2.** Seven representative AlphaFold cavities that are similar to the zafirlukast binding site in cysteinyl leukotriene receptor 1.

| **UniProt ID** | **Protein Name** | **Cavity ID** | **PMSmax** | **Vina Score (kcal/mol)** |
| --- | --- | --- | --- | --- |
| Q8TDU9 | Relaxin-3 receptor 2 | 1 | 0.849 | -9.80 |
| Q9UL12 | Sarcosine dehydrogenase, mitochondrial | 5 | 0.835 | -10.81 |
| Q96S06 | Lipase maturation factor 1 | 1 | 0.826 | -9.35 |
| Q12887 | Protoheme IX farnesyltransferase, mitochondrial | 2 | 0.822 | -10.60 |
| P06133 | UDP-glucuronosyltransferase 2B4 | 1 | 0.810 | -12.15 |
| Q9H568 | Actin-like protein 8 | 1 | 0.806 | -11.56 |
| Q9H270 | Vacuolar protein sorting-associated protein 11 homolog | 3 | 0.805 | -10.31 |

## Notes

### Competing Interest Statement

The authors have declared no competing interest.

